## Supplemental Figures and Tables for "A mutational atlas for Parkin proteostasis"

*Supplemental material*

**Fig. S1**, Low level of endogenous Parkin in HEK293T cells. p.2
**Fig. S2**, Correlations between independent repeats. p.3
**Fig. S3**, Correlations with the relative abundance of Parkin variants in U2OS cells. p.4
**Fig. S4**, Surface exposed beta strands display alternating mutational sensitivity. p.5
**Fig. S5**, $Zn^{2+}$ coordinating residues are sensitive to mutations. p.6
**Fig. S6**, High abundance variants are positioned in exposed regions. p.7
**Fig. S7**, Stabilization of Parkin by mutation of the active site. p.8
**Fig. S8**, Hyperactive Parkin variants display a reduced abundance. p.9
**Fig. S9**, Low stability tiles are buried in the Parkin structure. p.10
**Fig. S10**, Side-by-side comparison of the abundance map with the computational maps. p.11
**Fig. S11**, Correlation with Parkin melting temperatures. p.12
**Fig. S12**, Positioning of the pathogenic variants in the Parkin structure. p.13

**Table S1**, Data for the pathogenic and benign variants included in this study. p.14
**Table S2**, Primers used in this study. p.15

**References for supplemental material** p.16

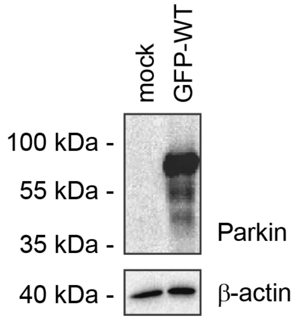


**Figure S1** – *Low level of endogenous Parkin in HEK293T cells.*
The level of Parkin in the HEK293T landing pad cell line was analyzed by SDS-PAGE and western blotting using antibodies to Parkin. β-actin served as a loading control. Note that we were unable to detect any endogenous Parkin in the cells.

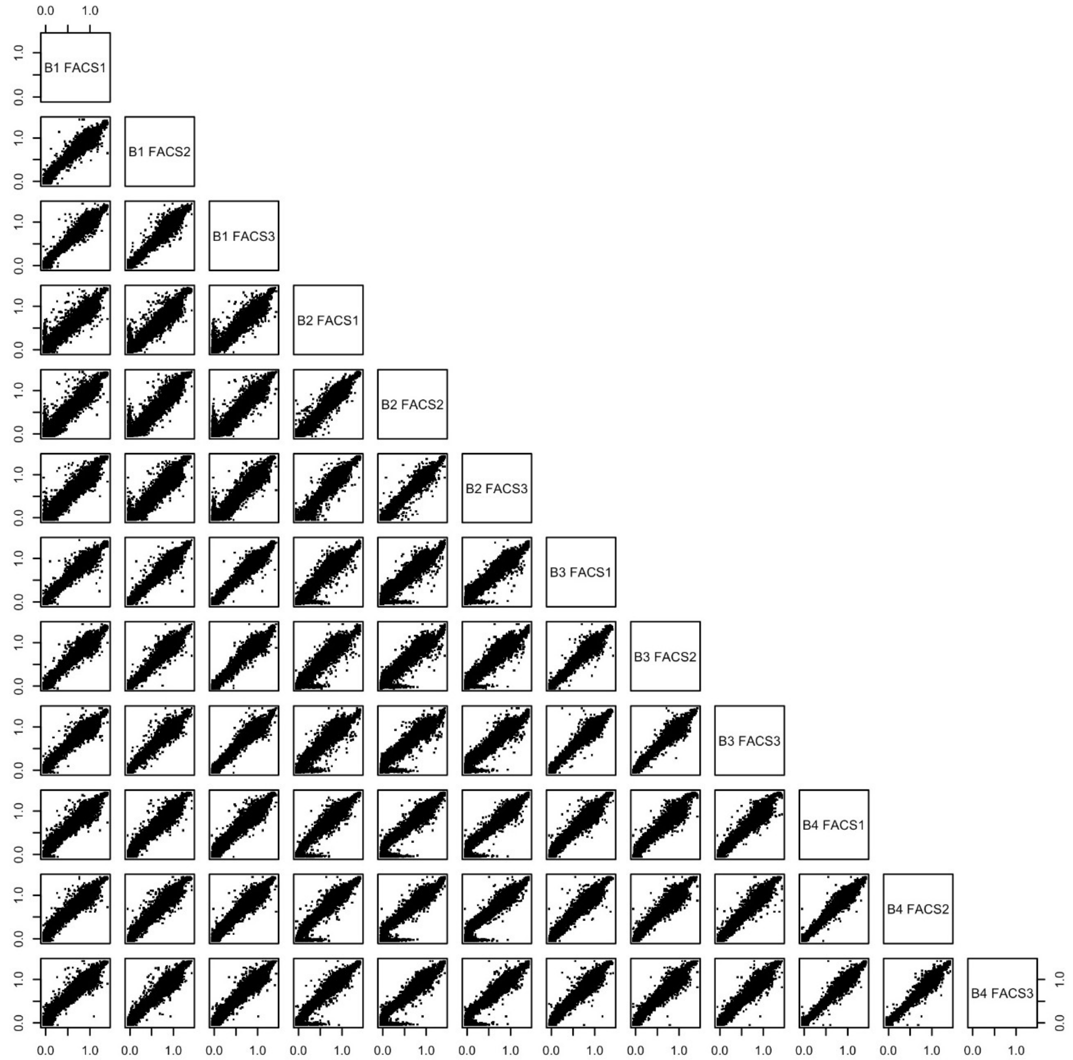


**Figure S2** – *Correlations between independent repeats.*

Reproduction of single amino acid variant scores between all four biological replicates (B1-B4) each with three FACS replicates. All Pearson correlations are in the range 0.96 to 0.99.

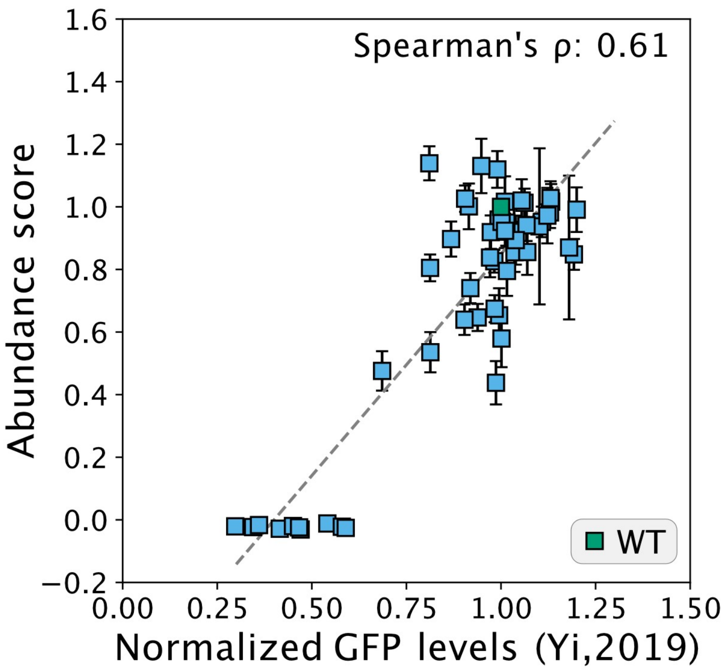


**Figure S3** – *Correlations with the relative abundance of Parkin variants in U2OS cells.*

Correlation of Parkin variant abundance with their relative abundance in U2OS cells based on fluorescence microscopy (Yi et al., 2019). The error bars indicate the standard deviation.

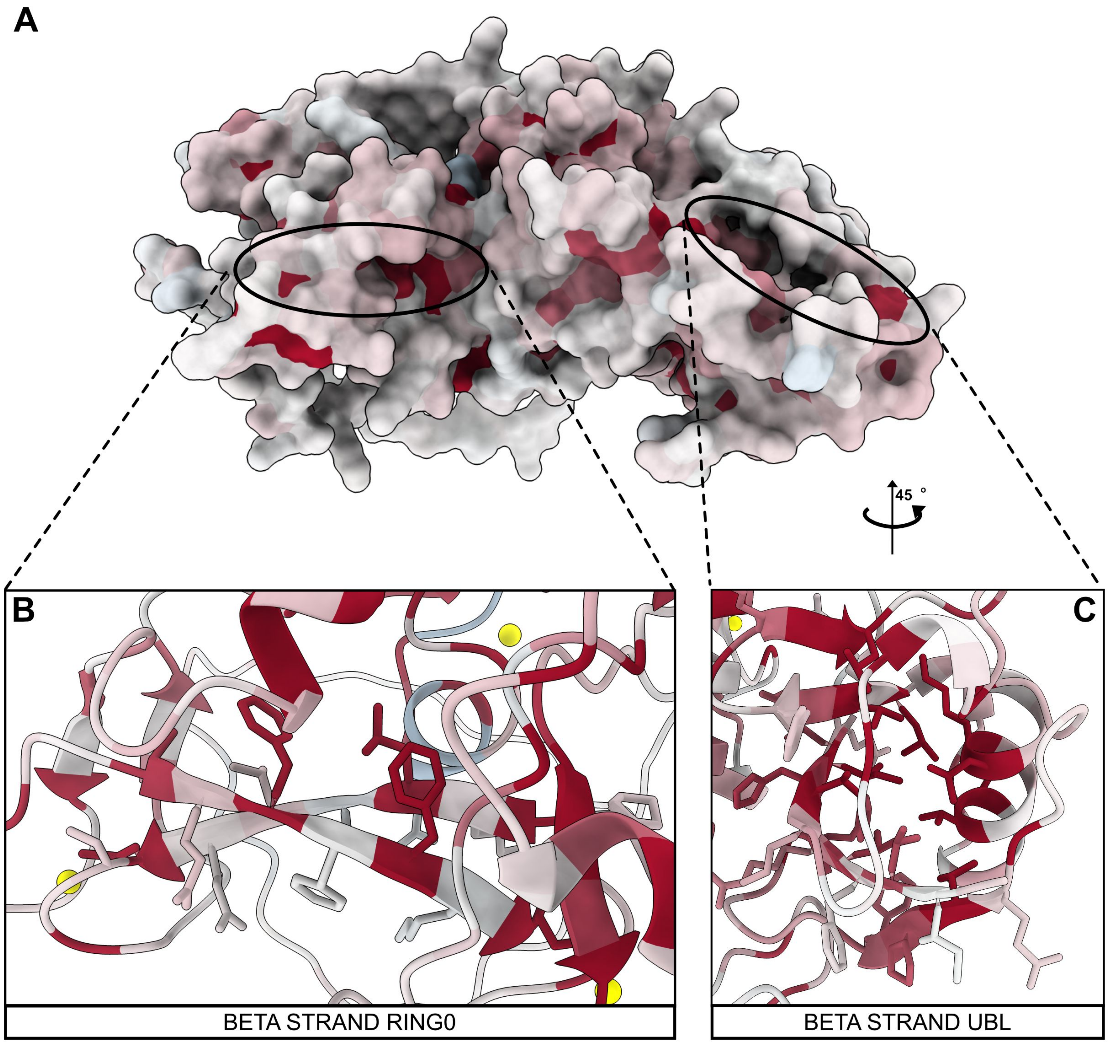


**Figure S4** – *Surface exposed beta strands display alternating mutational sensitivity.*
Examples of the alternating mutational patterning of the β-strands are shown for the RING0 (AB) and UBL (AC) domains. The structure is colored based on the median abundance score per residue.

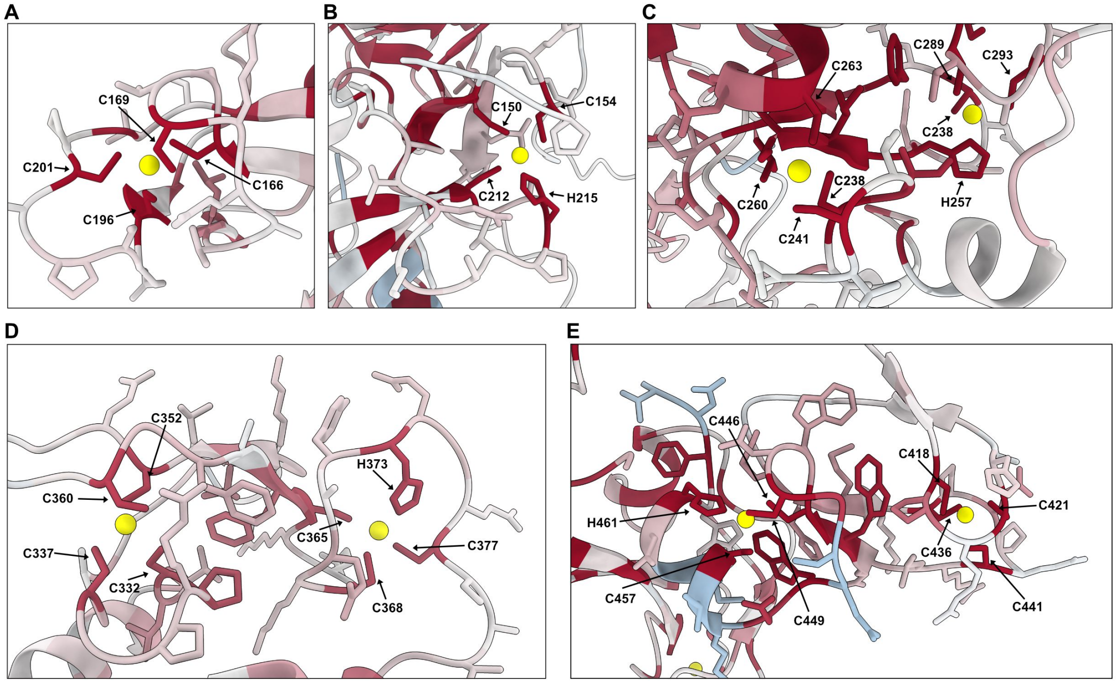


**Figure S5** – *$Zn^{2+}$ coordinating residues are sensitive to mutations.*
Zoom-in views on the $Zn^{2+}$ binding residues in the Parkin structure in the (AB) RING0 domain, (C) the RING1 domain, (D) IBR domain and (E) RING2 domain. The structure is colored based on the median abundance score per residue.

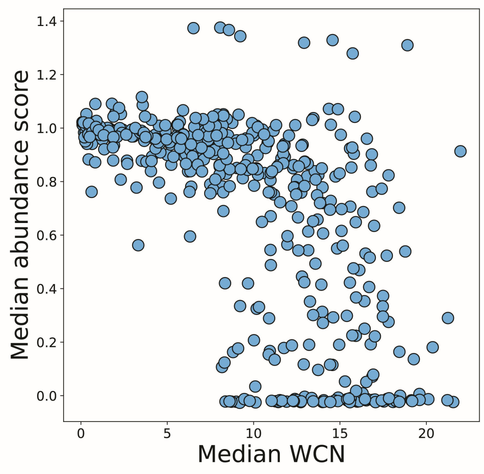


**Figure S6 -** *High abundance variants are positioned in exposed regions.*
Plot of the median abundance score per position and the weighted contact number (WCN).

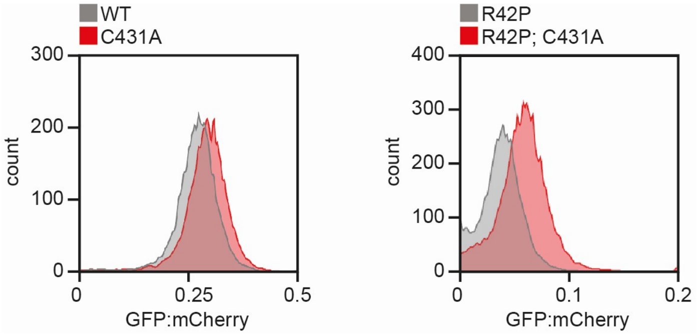


**Figure S7** – *Stabilization of Parkin by mutation of the active site.*

Flow cytometry profiles displaying wild-type (WT), catalytically dead (C431A), low abundance (R42P), and double mutant (R42P C431A) Parkin variants. Note that the R42P C431A double mutant displays a slightly increased level compared to the R42P single variant. Note that the scale on the x-axis is linear and the stabilization conferred by the C431A mutation is therefore minor.

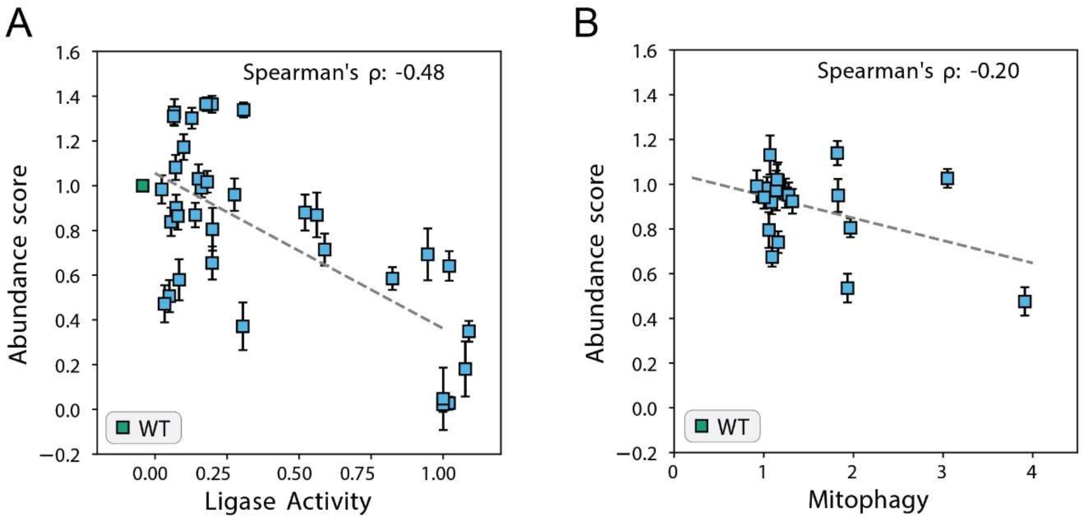


**Figure S8** – *Hyperactive Parkin variants display a reduced abundance.*
(A) Correlation of Parkin abundance scores with the relative activity of hyperactive Parkin variants evaluated by *in vitro* ubiquitination assays (Stevens et al., 2023). (B) Correlation of Parkin abundance scores with the relative CCCP-induced mitophagy activity of hyperactive Parkin variants normalized to their abundance (Yi et al., 2019). Error bars indicate the standard deviation.

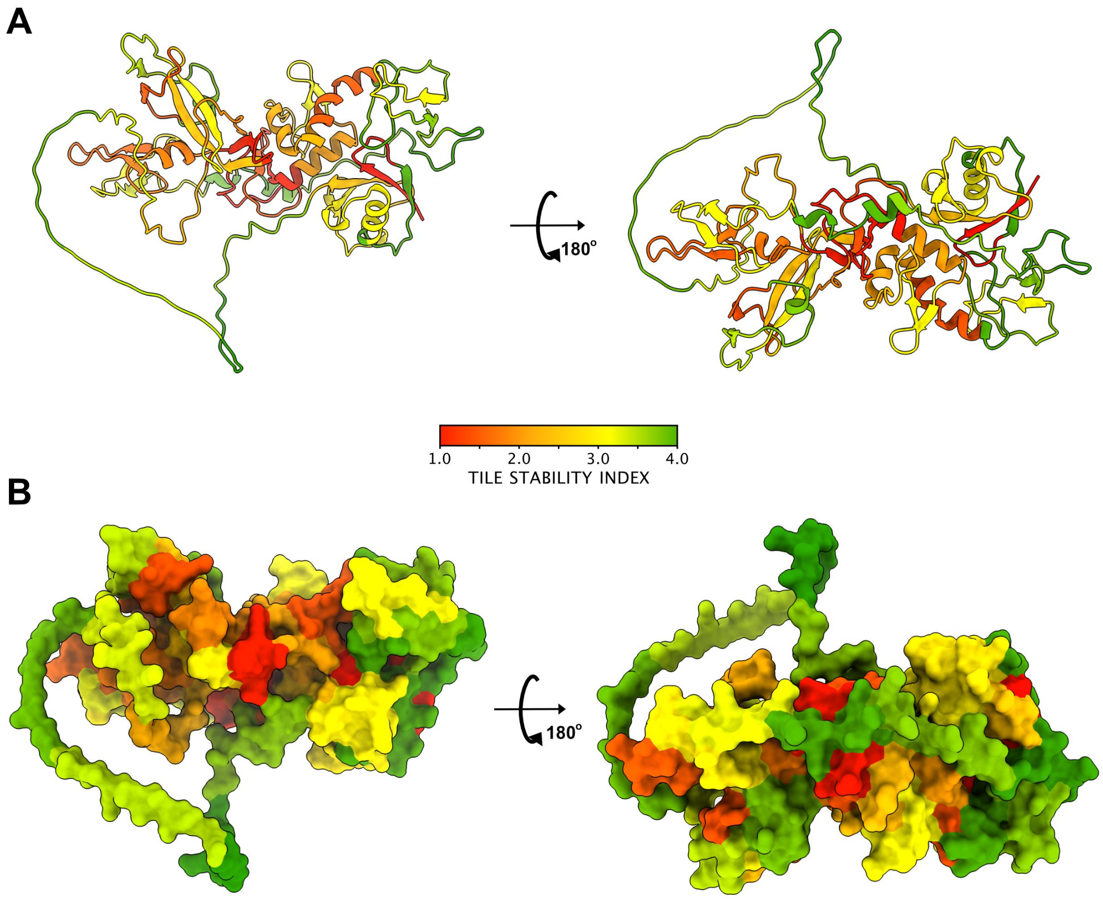


**Figure S9** – *Low stability tiles are buried in the Parkin structure.*
(A) Cartoon and (B) space filling representations of the Parkin structure colored by TSI. Note that regions with low stability are mostly buried in the structure.

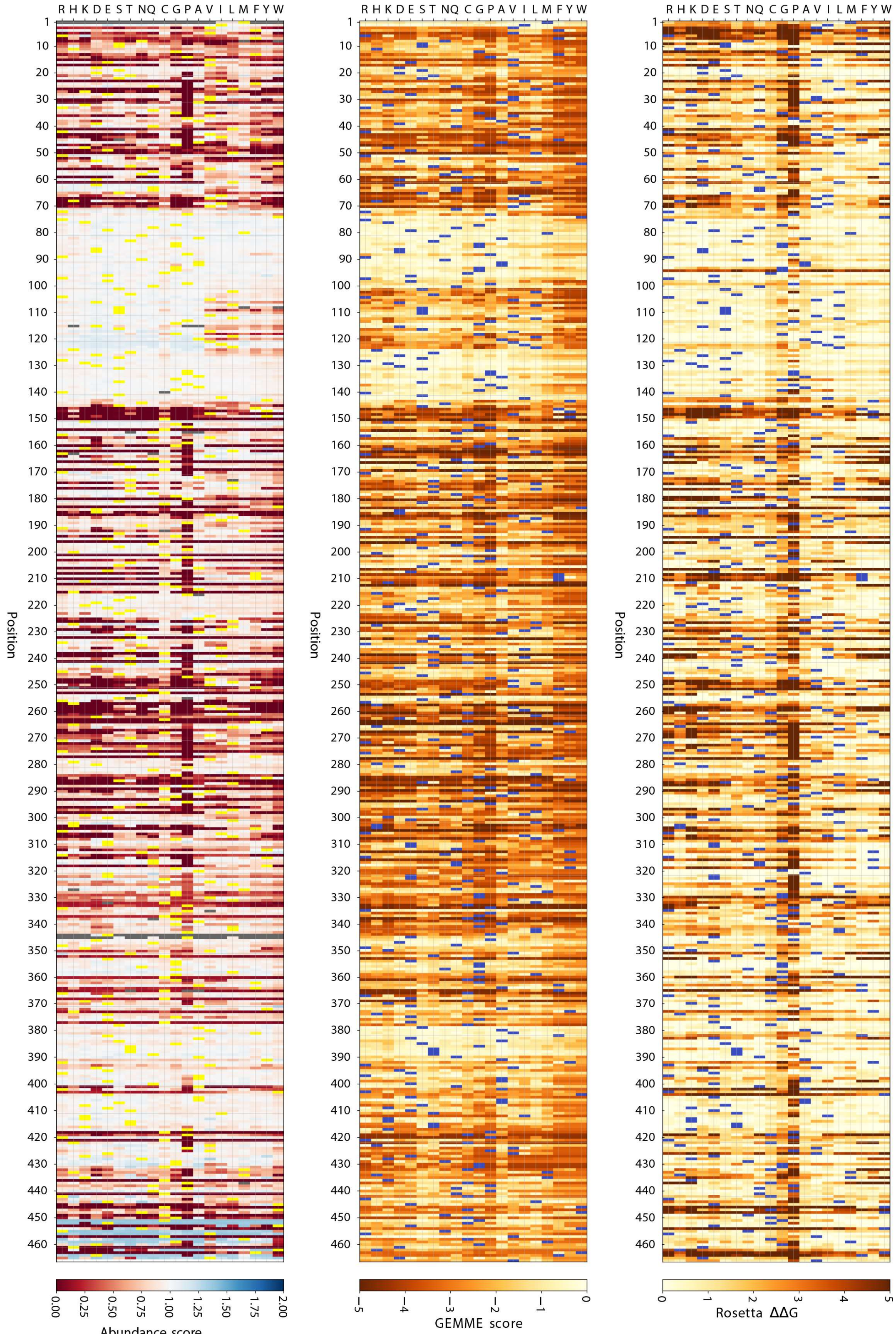


**Figure S10** - *Side-by-side comparison of the abundance map with the computational maps.*
The figure depicts the abundance map as determined by VAMP seq (left) next to the *in silico* maps based on evolutionary conservation by GEMME (middle) and structural stability (ΔΔG) determined by Rosetta (right).

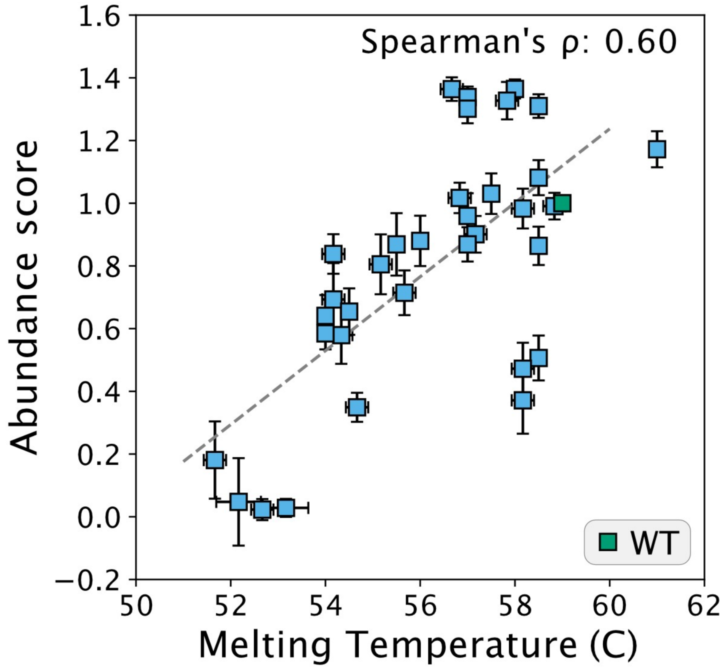


**Figure S11** – *Correlation with Parkin melting temperatures.*

Correlation of Parkin variant abundance scores with their corresponding melting temperatures determined by Stevens *et al*. (Stevens et al., 2023).

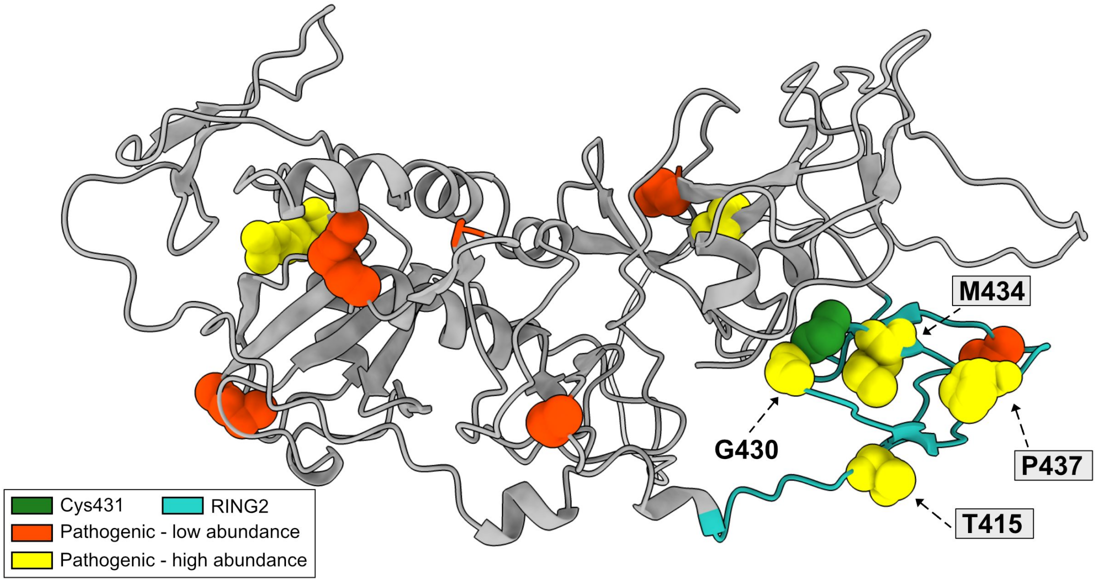


**Figure S12** – *Positioning of the pathogenic variants in the Parkin structure.*
Cartoon representation of the Parkin structure with pathogenic low abundance (red) and high abundance (yellow) variants marked, along with the active site cysteine at position 431 (green). The catalytic RING2 domain is marked in cyan. Note that several of the pathogenic high abundance variants cluster near the active site in the RING2 domain.

**Table S1**

*Data for the pathogenic and benign variants included in this study*

| Variant position | VAMP score | GEMME score | Rosetta ΔΔG score | gnomAD allele freq. | ClinVar category | Notes |
|---|---|---|---|---|---|---|
| Wild-type | 1 | 0 | 0 | - | Benign | - |
| D18N | 0.99 | -0.09 | 0.51 | 3.40E-04 | Benign | - |
| R33Q | 0.84 | -0.41 | 0.23 | 7.85E-05 | Pathogenic | - |
| Q34R | 0.90 | -1.66 | 0.90 | 3.05E-03 | Benign | - |
| R42C | 1.02 | -1.46 | 0.56 | 1.47E-04 | Benign | - |
| R42P | -0.02 | -3.65 | 3.85 | 2.53E-05 | Pathogenic | - |
| A46T | 0.65 | -1.58 | 2.12 | 3.17E-03 | Benign | - |
| A82E | 1.03 | -0.15 | -0.50 | 3.06E-03 | Conflicting | Reclassified, benign* |
| P153R | 0.98 | -1.98 | 1.11 | 2.77E-03 | Benign | - |
| K161N | 0.94 | -3.98 | 1.50 | 6.98E-06 | Pathogenic | - |
| S167N | 0.95 | -0.33 | 0.66 | 0.06 | Benign | - |
| M192L | 1.02 | -0.93 | -0.09 | 0.01 | Benign | - |
| C212Y | -0.01 | -5.27 | 6.41 | 1.19E-05 | Pathogenic | - |
| V248I | 0.92 | -1.40 | 0.55 | 1.17E-04 | Benign | - |
| C253Y | -0.03 | -4.55 | 2.61 | 1.59E-05 | Pathogenic | - |
| L272I | 0.91 | -1.77 | 0.78 | 7.60E-05 | Benign | - |
| R275W | -0.02 | -3.57 | 1.73 | 2.02E-03 | Pathogenic | - |
| G284R | 0.44 | -4.85 | 2.00 | 1.19E-05 | Pathogenic | - |
| R334C | 0.48 | -0.87 | 0.75 | 1.72E-03 | Conflicting | Reclassified, benign* |
| R366W | 0.74 | -4.03 | 0.57 | 2.23E-04 | Benign | - |
| V380L | 0.92 | -0.06 | 0.44 | 0.17 | Benign | - |
| D394N | 0.85 | -0.25 | 0.14 | 0.03 | Benign | - |
| R402C | 0.96 | -1.54 | 1.38 | 1.91E-03 | Conflicting | Reclassified, benign* |
| T415N | 1.02 | -3.19 | 0.73 | 7.95E-06 | Pathogenic | - |
| G430D | 1.00 | -3.59 | -1.07 | 8.42E-05 | Pathogenic | - |
| M434T | 1.17 | -2.81 | 2.87 | # | Pathogenic | - |
| P437L | 0.87 | -1.12 | 0.77 | 1.80E-03 | Conflicting | Reclassified, pathogenic** |
| C441R | -0.02 | -3.40 | 1.61 | 5.20E-05 | Pathogenic | - |

#= variant not listed in gnomAD.

*= assigned new category based on Sherloc criteria (Nykamp et al., 2017; Yi et al., 2019).

**= assigned new category based on MDSgene database (Lill et al., 2016).

**Table S2**

*Primers used in this study*

| Name | Sequence |
|---|---|
| ASPA_PARK2_index2_re | ACGCAATTGCAGAACTAGTCCTCATATGTCCTGG |
| gDNA_2nd | AATGATACGGCGACCACCGAGATCTACAC(NNNNNNNN)CGTGACCGCCGCC |
| JS_R | CAAGCAGAAGACGGCATACGAGAT(NNNNNNNN)GGGTTAGCAAGTGGCAGCCT |
| LC1020 | CCAGGACATATGAGGACTAG |
| LC1031 | GGGTTAGCAAGTGGCAGCCTTCTCCTTAATCAGCTCTTCG |
| LC1040 | AAGAACCGCTAGAAGCGTCGCTGTACAAATAGTT |
| LC1041 | CGAGAAAGCTAGCGCAAACGACTACTCGCA |
| LC1042 | CTGATTAAGGAGAAGGCTGCCACTTGCTAACCC |
| PCR2_Fw | AATGATACGGCGACCACCGAGATCTACAC(NNNNNNNN)CCAGGACATATGAGGACTAG |
| VV1 | TAGTAACTTAAGAATTCACCGGTCTGACCT |
| VV2 | GGTGGCTCCGCTGCCTCTA |
| VV2S | GGGTTAGCAAGTGGCAGCCTTGCAGACCTGAAGAGAGGGA |
| VV3 | AAGACGCGTTCTAGAGGCAGC |
| VV4 | AGGTCAGACCGGTGAATTCTTAAGTTACTA |
| VV16 | CGGTCACGAACTCCAGCAGGACCATGTG |
| VV18 | GGGAGAAGGAGGTCAGACCGGTGAATTCTTAAGTTACTA |
| VV19 | TCTGCAAGGCTGCCACTTGCTAACCC |
| VV21 | GAGTGATCCCGGCGGCGGTCACG |
| VV40S | GAGAACGTATGTCGAGGTAGGC |
